# Global protein responses of multi-drug resistant plasmid containing *Escherichia coli* to ampicillin, cefotaxime, imipenem and ciprofloxacin

**DOI:** 10.1101/2021.07.26.453744

**Authors:** Anatte Margalit, James C. Carolan, Fiona Walsh

## Abstract

Antimicrobial resistance (AMR) and multi-drug resistance (MDR) in pathogenic bacteria are frequently mediated by plasmids. However, plasmids do not exist in isolation but rather require the bacterial host interaction in order to produce the AMR phenotype. This study aimed to utilise mass spectrometry-based proteomics to reveal the plasmid and chromosomally derived protein profile of *Escherichia coli* under antimicrobial stress. This was achieved by comparing the proteomes of *E. coli* containing the MDR pEK499 plasmid, under ampicillin, cefotaxime, imipenem or ciprofloxacin stress with the proteomes of these bacteria grown in the absence of antimicrobial. Our analysis identified statistically significant differentially abundant proteins common to groups exposed to the β-lactam antimicrobials but not ciprofloxacin, indicating a β-lactam stress response to exposure from this class of drugs, irrespective of β-lactam resistance or susceptibility. These include ecotin and free methionine-R-sulfoxide reductase. These data also identified distinct differences in the cellular response to each β-lactam. Data arising from comparisons of the proteomes of ciprofloxacin-treated *E. coli* and controls detected an increase in the relative abundance of proteins associated with ribosomes, translation, the TCA-cycle and several proteins associated with detoxification and a decrease in the relative abundances of proteins associated with stress response, including oxidative stress. We identified changes in proteins associated with persister formation in the presence of ciprofloxacin but not the β-lactams. The plasmid proteome differed across each treatment and did not follow the pattern of antimicrobial – AMR protein associations. For example, a relative increase in the amount of blaCTX-M-15 in the presence of cefotaxime and ciprofloxacin but not the other β-lactams, suggesting regulation of the blaCTX-M-15 protein production. The proteomic data from the this study provided novel insights into the proteins produced from the chromosome and plasmid under different antimicrobial stresses. These data also identified novel proteins not previously associated with AMR or antimicrobials responses in pathogens, which may well represent potential targets of AMR inhibition.

## Introduction

Multi-drug resistance (MDR) plasmids are reducing the effectiveness of our antimicrobial arsenal in a wide range of pathogens globally. The World Health Organisation (WHO) recognised that extended spectrum beta-lactamase (ESBL) *Escherichia coli* are on the critical list of priority pathogens in relation to human health (1). The extended spectrum ß-lactamases (ESBLs) are most frequently disseminated within and across pathogen species via horizontal transfer of plasmids. The response of pathogens such as *E. coli* to antimicrobials is most commonly measured via antimicrobial susceptibility testing for phenotypic detection of resistant bacteria followed by genotypic or genomic identification of the resistance mechanisms. The focus of such analysis is to identify the treatment options available and understand the mechanisms of resistance. The pathogen response to the antimicrobial is measured in relation to its classification as resistant or susceptible and the specific resistance gene present e.g. an ESBL is identified via testing the bacterial response to ß -lactams and their inhibitors and followed by screens for a variety of ESBL genes. Clinical laboratories and pathogen studies are starting to use whole genome sequencing as a high-throughput method for antimicrobial resistance (AMR) detection. However, as with all organisms, the bacterial genotype does not necessarily directly dictate the phenotype as numerous regulatory mechanisms are also involved.

Antimicrobial resistance mechanisms are frequently plasmid mediated and many of the plasmids confer resistance to several antimicrobials concurrently. One such pathogen - plasmid combination is the internationally prevalent *E. coli* O25:H4-ST131 containing the plasmid pEK499 (117,536 bp in size). The pEK499 plasmid harbours the 10 antimicrobial resistance genes: *bla*CTX-M-15, *bla*OXA-1, *bla*TEM-1, *aac6’-Ib-cr, mph(A), catB4, tet(A)*, integron-borne *dfrA7, aadA5, sulI* genes and several hypothetical and conjugation associated genes (2). This plasmid confers resistance to beta-lactams, macrolides, chloramphenicol, tetracycline, trimethoprim, streptomycin, spectinomycin and sulphonamide and reduced susceptibility to ciprofloxacin. In addition, the clonal expansion of *E. coli* with the sequence type 131 has significantly increased the global dissemination of *bla*CTX-M-15.

We know relatively little about the entire bacterial-plasmid system response of pathogens to antimicrobial stress. Plasmids must interact with the bacterial cell in order to transcribe and translate their DNA into proteins and these proteins must interact with bacterial proteins in order to be exported or illicit change resulting in resistance. Although interactions between the plasmid and the bacterial host clearly exist, the study of plasmid mediated AMR focus mainly on the plasmid mediated genes or phenotypic response of the host pathogen itself. Therefore the identification and characterisation of the pathways or proteins required for the bacteria to produce proteins associated with AMR could provide novel targets to restrict AMR.

A recent study provided an insight into how proteomics may be used in an unbiased fashion for the detection of AMR in pathogenic bacteria cultured in the absence of antimicrobial (3). The authors followed a workflow that resulted in 98% sensitivity and 100% specificity across seven pathogens and 11 AMR determinants, thus demonstrating the applicability of such proteomic workflows in clinical microbiology. If such a system is implemented then the additional data could prove valuable in understanding the complex plasmid – bacterial host interactions required for the production of the AMR proteins. Few studies currently exist on the proteomic response of pathogens containing AMR plasmids to antimicrobials. These include adaptation of *bla*CTX-M-1 containing *E. coli* to cefotaxime (4) and the global response of tetracycline resistant *E. coli* to oxytetracycline (5). We have not identified any publication analysing the proteomic changes of MDR plasmid mediated resistance under different antimicrobial stresses, such as the study presented here.

Our study aimed to analyse the responses of the clonal *E. coli* ST131 strain, containing pEK499 plasmid under different antimicrobial stresses and compare them with the proteome in the absence of antimicrobial stresses in order to identify common and antimicrobial specific global response of the *E. coli* and plasmid proteomes to antimicrobial stress. The aim was to specifically understand how the plasmid and bacterial host proteins were influenced by the different antimicrobial stresses. By analysing these factors systematically we aimed to identify the pathways and antimicrobial specific responses of the bacteria. Using proteomics we provide an unbiased protein map of a pathogen with a MDR plasmid under antimicrobial stresses.

## Materials and Methods

### Preparation of *Escherichia coli* proteins for mass spectrometry

The bacterial strain *Escherichia coli* NCTC 13400, containing the MDR conjugative plasmid pEK499, was used in all experiments. The pEK499 plasmid was 117,536 bp in length and belongs to incompatibility group F as represented a fusion of two replicons of types FII and FIA (2). *Escherichia coli* (NCTC 13400) containing the MDR plasmid pEK499 was exposed to antimicrobials for which the bacteria displayed a resistance phenotype (ampicillin 64mg/L, cefotaxime 256 mg/L) and those, which there was no resistance phenotype (imipenem 0.06 mg/L, ciprofloxacin 0.06 mg/L) (2). The control comprised the *E. coli* with pEK499 grown without antimicrobial. All strains were grown separately in Luria-Bertani (LB) at 37 °C with shaking at 200 rpm. All experiments were performed in biological triplicates. Cells were harvested by centrifugation at 3000 rpm for 15 minutes. The cell pellet was resuspended in ammonium bicarbonate (1 ml, 50 Mm, pH 7.8) and sonicated on ice in 10 second bursts five times. The lysate was subjected to centrifugation at 13,000 rpm to collect the cellular debris. The supernatant was quantified using the Qubit™ quantification system (Invitrogen), following the manufacturer’s instructions. The protein sample was reduced by adding 5 μl 0.2 M dithiothreitol (DTT) and incubated at 95°C for 10 minutes, followed by alkylation with 0.55 M iodoacetamide (4 µl) at room temperature, in the dark for 45 minutes. Alkylation was stopped by adding DTT (20 µl, 0.2 M) and incubation for 45 minutes at 25 °C. Sequence Grade Trypsin (Promega) (0.5 µg/µl) was added to the proteins and incubated at 37°C for 18 hours. The digested protein sample was brought to dryness using a Speedyvac concentrator (Thermo Scientific Savant DNA120). Samples were purified for mass spectrometry using C18 Spin Columns (Pierce), following the manufacturer’s instructions. The eluted peptides were dried in a SpeedyVac concentrator (Thermo Scientific Savant DNA120) and resuspended in 2% *v/v* acetonitrile and 0.05% *v/v* Trifluoroacetic acid (TFA) to give a final peptide concentration of 1 µg/µl. The samples were sonicated for five minutes to aid peptide resuspension, followed by centrifugation for five minutes at 13,000 rpm. The supernatant was removed and used for mass spectrometry. Three independent biological replicates for each group were analysed.

### Mass Spectrometry: LC/MS Xcalibur Instrument parameters for proteomic data acquisition

Digested proteins (1 µg) isolated from the replicates for each *E. coli* sample were loaded onto a QExactive (ThermoFisher Scientific) high-resolution accurate mass spectrometer connected to a Dionex Ultimate 3000 (RSLCnano) chromatography system. Peptides were separated by an increasing acetonitrile gradient on a 50 cm EASY-Spray PepMap C18 column with 75 µm diameter (2 µm particle size), using a 180 minute reverse phase gradient at a flow rate of 300 nL/mi ^-1^ n. All data were acquired over 141 minutes, with the mass spectrometer operating in an automatic dependent switching mode. A full MS scan at 140,000 resolution and a range of 300 – 1700 *m/z*, was followed by an MS/MS scan at 17,500 resolution, with a range of 200-2000 *m/z* to select the 15 most intense ions prior to MS/MS.

Quantitative analysis (protein quantification and LFQ normalization of the MS/MS data) of the *E. coli* proteome arising from exposure to the different antimicrobials, was performed using MaxQuant version 1.6.3.3 (http://www.maxquant.org) following the general procedures and settings outlined in Hubner et al., 2010 (6). The Andromeda search algorithm incorporated in the MaxQuant software was used to correlate MS/MS data against the Uniprot-SWISS-PROT database for *E. coli* K12 (4319 entries) and the *E. coli* strain plasmid pEK499 (141 entries). The following search parameters were used: first search peptide tolerance of 20 ppm, second search peptide tolerance 4.5 ppm with cysteine carbamidomethylation as a fixed modification and N-acetylation of protein and oxidation of methionine as variable modifications and a maximum of two missed cleavage sites allowed. False discovery rate (FDR) was set to 1 % for both peptides and proteins, and the FDR was estimated following searches against a target-decoy database. Peptides with a minimum length of seven amino acid length were considered for identification and proteins were only considered identified when observed in three replicates of one sample group.

### Data Analysis of the proteome

Perseus v.1.5.5.3 (www.maxquant.org/) was used for data analysis, processing and visualisation. Normalised LFQ intensity values were used as the quantitative measurement of protein abundance for subsequent analysis. The data matrix was first filtered for the removal of contaminants and peptides identified by site. LFQ intensity values were log_2_ transformed and each sample was assigned to its corresponding group. Proteins not found in all three replicates in at least one group were omitted from the analysis. A data-imputation step was conducted to replace missing values with values that simulate signals of low abundant proteins chosen randomly from a distribution specified by a downshift of 1.8 times the mean standard deviation (SD) of all measured values and a width of 0.3 times this SD.

Normalised intensity values were used for a principal component analysis (PCA). Exclusively expressed proteins (those that were uniquely expressed or completely absent in one group) were identified from the pre-imputation dataset (Supplemental dataset 1) and included in subsequent post-imputation analyses (Supplemental dataset 2). To visualise differences between two samples, pairwise Student’s t-tests were performed for all using a cut-off of p<0.05 on the post-imputated dataset. Volcano plots were generated in Perseus by plotting negative log p-values on the y-axis and log_2_ fold-change values on the x-axis for each pairwise comparison. The ‘categories’ function in Perseus was utilized to highlight and visualise the distribution of various pathways and processes on selected volcano plots. Statistically significant (ANOVA, p<0.05) proteins were chosen for further analysis. Gene ontology (GO) mapping was also performed in Perseus using the UniProt gene ID for all identified proteins to query the Perseus annotation file (downloaded September 2018) and extract terms for gene ontology biological process (GOBP), gene ontology cellular component (GOCC), gene ontology molecular function (GOMF) and Kyoto Encyclopedia of Genes and Genomes (KEGG) name. Enrichment analysis was performed in Search Tool for the Retrieval of Interacting Genes/Proteins (STRING), using a high confidence setting (0.700), and hiding disconnected nodes in the network. Statistically significant protein names arising from pairwise t-tests were inputted into the STRING database to identify interactions occurring between proteins that were increased or decreased in relative abundance between a treatment and the control. The MS proteomics data and MaxQuant search output files have been deposited to the ProteomeXchange Consortium (7) via the PRIDE partner repository with the dataset identifier PXD027164.

## Results

Label free quantitative (LFQ) proteomics was employed to investigate the proteomic response of *E. coli* pEK499 when exposed to different antimicrobials. The pEK499 plasmid confers resistance to ampicillin and cefotaxime, both of which were added to the bacterial culture above the break-point (the concentration of antimicrobial used to define whether an infection by the pathogenic species is likely to be treatable in a patient). This strain is susceptible to imipenem and ciprofloxacin and was exposed to these antibiotics at sub-minimum inhibitory concentration (MIC) levels (2). In total, 1586 proteins were initially identified, of which 945 (16 of which were of plasmid origin; Supplemental dataset 7) remained after filtering and processing (Supplemental dataset 2). The PCA performed on all filtered proteins, resolved only the ciprofloxacin treated *E. coli* and separated those samples from all other samples along component 1 (Fig. 1). Principle components 1 and 2 accounted for 40.6 % of the total variance within the data. The samples obtained from bacteria exposed to cell wall biosynthesis inhibitors ampicillin, cefotaxime and imipenem grouped close to the control, thereby indicating fewer changes to the proteome in bacteria exposed to this group of antimicrobials compared to the control relative to the replicates exposed to ciprofloxacin. The ciprofloxacin exposed samples were furthest from the control, indicating a significant change to the protein profile in this group compared to the control.

**Figure 1.**
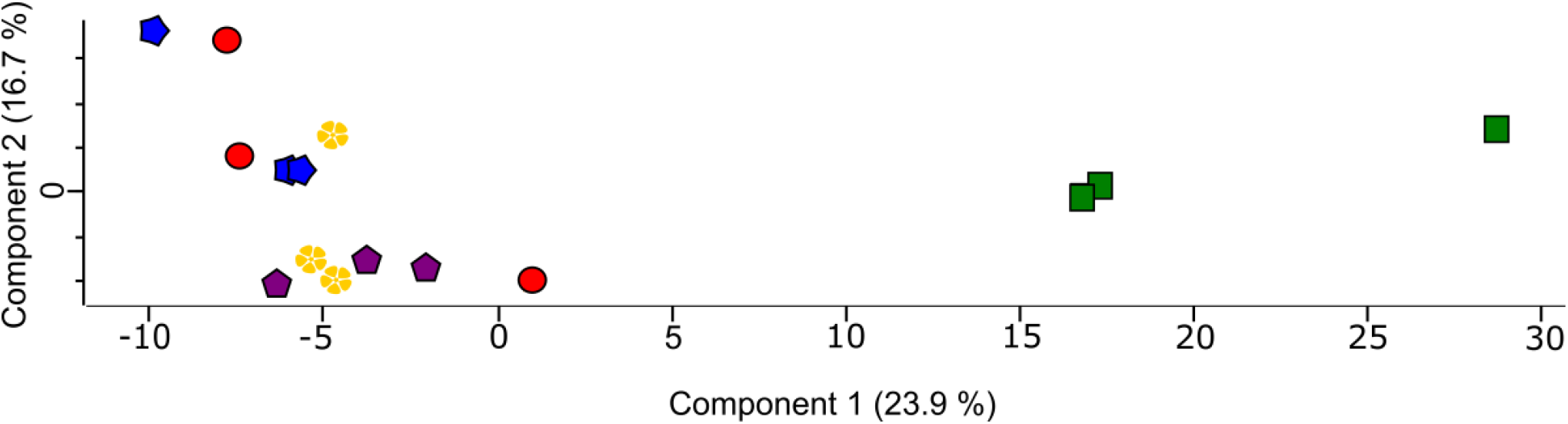
Principal component analysis of the proteomes of pEK499 containing *E. coli* treated with ampicillin (red), cefotaxime (blue), imipenem (yellow) or ciprofloxacin (green) and the control untreated bacteria (purple).

Volcano plots were produced by pairwise Student’s t-tests (p <0.05) on the post-imputated dataset to determine differences in protein abundance between two groups (Fig. 2A-D). Statistically significant differentially abundant (SSDA) proteins arising from pairwise t-tests were determined between the groups and included 95 for ampicillin vs control, 145 for cefotaxime vs control, 89 for imipenem vs control and 208 ciprofloxacin vs control (Supplemental dataset 3-6). The 20 most differentially abundant proteins between each group are highlighted and labelled on the volcano plots (Fig. 2A-D).

**Fig. 2.**
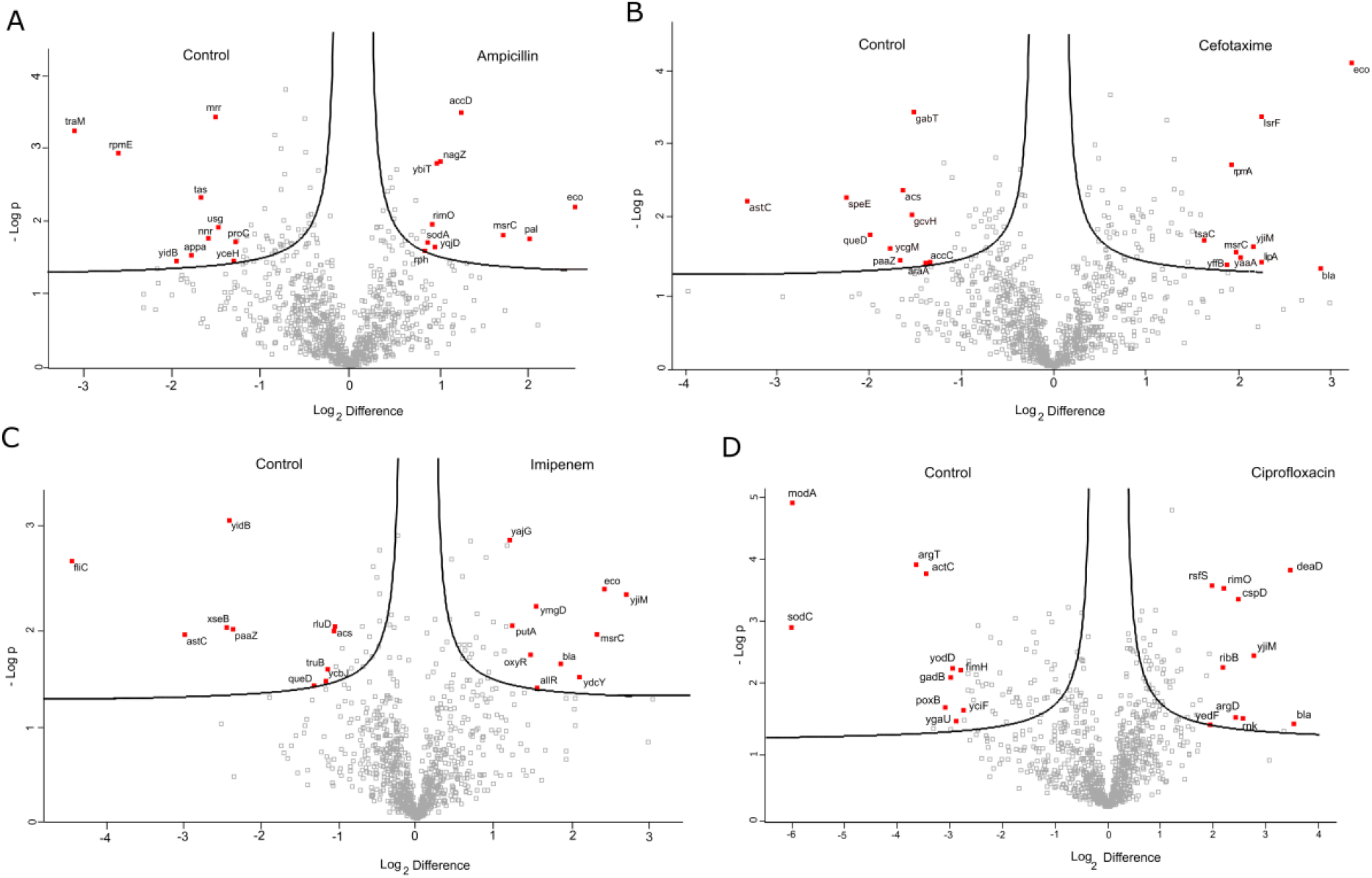
A-D Volcano plots derived from pairwise comparisons between A) *E. coli* pEK499 treated with ampicillin and control, B) cefotaxime and control, C) imipenem and control and D) ciprofloxacin and control. The distribution of quantified proteins according to p value (−log10 p-value) and fold change (log2 mean LFQ intensity difference) are shown. Proteins above the line are considered statistically significant (p-value <0.05). The top 20 most differentially abundant proteins are shown for each group.

The proteomic data arising from pairwise t-tests revealed an increase in the relative abundance of several proteins common to groups exposed to ampicillin, cefotaxime and imipenem compared with the control (Supplemental datasets 3 – 6). Among these were stress-related proteins including ecotin (Eco), and methionine-R-sulfoxide reductase (MsrC). Compared to the controls, a statistical difference in the relative abundances of two beta-lactamases blaCTX-M-15 and blaTEM-1 was detected in the cefotaxime and imipenem-exposed groups, respectively, but not ampicillin-exposed bacteria. Additionally, the relative abundance of proteins involved with detoxification were increased in bacteria treated with these cell-wall inhibitors, including superoxide dismutase SodA (ampicillin- and cefotaxime-treated), peroxiredoxin OsmC, glutaredoxin 3 (GrxC) and 4 (GrxD) (cefotaxime-treated), hydrogen peroxide-inducible genes activator (OxyR) (imipenem treated) and thiosulfate sulfurtransferase PspE (cefotaxime- and imipenem-treated) (8 - 11). Cold shock proteins (CspE and CspA in cefotaxime treated and CspE in imipenem-treated groups) were detected at higher levels in these groups compared to the controls and ampicillin-treated bacteria.

Differential changes in the abundance of cell wall biosynthesis proteins were detected in each group exposed to cell-wall inhibitors including β-hexosaminidase NagZ (ampicillin- and cefotaxime-treated), cell division coordinator CpoB (cefotaxime and imipenem-treated), UDP-N-acetylmuramoylalanine--D-glutamate ligase MurD (imipenem-treated) and peptidoglycan-associated lipoprotein Pal (ampicillin-treated). Alanine racemase (DadX) was detected at lower levels in cefotaxime-treated cells compared to the other groups and the control. There were changes in the relative abundance of proteins involved in cell division processes amongst all groups treated with cell wall inhibitors compared to the control. These included increases in the cell division protein FtsZ observed in the cefotaxime- and imipenem-treated groups (1.50-fold increase and 1.16-fold increase, respectively). However, the negative modulator of initiation of replication SeqA was increased in the ampicillin-treated samples (1.61-fold).

The categories of proteins with decreased relative abundances, were quite dissimilar between the groups treated with cell wall inhibitors. For example, in the ampicillin-treated groups, proteins with the greatest decrease in relative abundance were associated with the uptake of foreign DNA and DNA processing (Relaxosome protein TraM; 8.61-fold decrease, YidB; 3.73-fold decrease, Mrr restriction system protein; 2.88-fold decrease) and in addition, macrolide resistance MphA (2.38-fold decrease). In the cefotaxime-treated groups, proteins with the greatest decrease in relative abundance were associated with amino acid metabolism. These included succinylornithine transaminase AstC (10-fold decrease) and polyamine aminopropyltransferase SpeE (4.74-fold decrease). In the imipenem-treated group, flagellin FliC protein was the most decreased in abundance (21.92-fold decrease), although proteins involved in amino acid metabolism were also decreased in abundance (succinylornithine transaminase AstC; 9.43-fold decrease, bifunctional protein PaaZ; 5.84-fold decrease). The outer membrane protein Slp was also reduced in the imipenem treated samples relative to the control.

Enrichment analysis of statistically significant proteins using STRING, identified differences in the protein pathways between antibiotic-treated groups and the controls, and provided insights into the protein-protein interactions that may be occurring within the groups. Analysis of the ampicillin-treated group and the control using STRING revealed a decrease in the relative abundance of several proteins associated with the ribosome and translation (Supplemental fig. 1B). In contrast, there was an increase in the relative abundance of proteins associated with these pathways in the cefotaxime-treated group (Supplemental fig. 2A), and interestingly, a decrease in the relative abundance of proteins involved in amino acid metabolism (Supplemental fig. 2B). Compared to the control, the relative abundance of proteins involved with carbohydrate metabolism was decreased in the cefotaxime-and imipenem-treated groups (Supplemental fig. 2B and 3B). Specifically, the levels of proteins involved in the glycolytic pathway were reduced in the imipenem-treated groups (Supplemental fig. 3B).

Pairwise t-tests of the proteomic data arising from ciprofloxacin-treated *E. coli* and controls detected an increase in the relative abundance of proteins associated with ribosomes, translation, the TCA-cycle and several proteins associated with detoxification. A decrease in the relative abundance of proteins involved with glutathione metabolism and detoxification was also identified in the data set (Supplemental dataset 6), including acid stress chaperone HdeB (−6.04), periplasmic AppA protein (−5.63), peroxiredoxin OsmC (−5.20) and superoxide dismutase (−1.40). HdeB and OsmC are involved in the acid stress response in *E. coli*. Compared to the control, there was an increase in the relative abundance of several cold shock proteins (CspA, CspC, CspD and CspE). There was a general increase in the relative abundance of proteins associated with amino acid metabolism. Similar to the ampicillin-treated cells, the relative abundance of the relaxosome protein TraM, was decreased (6.62-fold decrease). The top three most differentially abundant proteins in this group were the plasmid mediated β-lactamase (blaCTX-M-15, 11.63-fold increase), ATP-dependent RNA helicase DeaD (11-fold increase) and YjiM, an uncharacterised protein (6.87-fold increase). The top three proteins with the greatest decrease in relative abundance included superoxide dismutase SodC (64.79-fold decrease), molybdate-binding periplasmic protein ModA (64.08-fold decrease) and lysine/arginine/ornithine-binding periplasmic protein ArgT (10.49-fold decrease). STRING analysis of the statistically significant protein set arising from comparisons between the ciprofloxacin-treated cells and the controls, highlighted the reduced levels of proteins associated with a stress response (Supplemental fig. 4B), and a distinct increase in the relative abundance of proteins associated with the translation and the ribosome, the Tricarboxylic acid (TCA) cycle and glycine, serine and threonine metabolism in the ciprofloxacin-treated group compared to the control (Supplemental fig. 4A).

### Variations across the pEK499 plasmid proteomes

The proteomes of the plasmids under antibiotic stress were analysed in a similar manner to the entire proteome, by comparison with the control proteome. The initial analysis was to identify the proteins detected with genetic origins to the plasmid. Post-imputation analysis revealed several changes in the relative abundance of proteins originating from the plasmid (Table 2). Only the addition of cefotaxime was associated with an increase in the protein of the corresponding resistance mechanism (blaCTX-M-15). Ampicillin and ciprofloxacin did not results in the increased protein abundances of any ß-lactamases or the Aac(6’)Ib-cr protein associated with reduced susceptibility to fluoroquinolones. However, these proteins were present in the control and thus it appears that this demonstrates that there is no additional regulation of their protein production in the presence of these antimicrobials. There were no carbapenemases present on the plasmid.

**Table 1:** pEK499 plasmid-derived proteins detected by mass spectrometry. Proteins encoded by genes present on the pEK499 plasmid were detected in all groups, or exclusive to specific groups of *Escherichia coli*.

| Protein ID | Protein name | Sample presence | Sample absence |
| --- | --- | --- | --- |
| ACQ41977.1 | Orf1176 protein (SopA) | All bacterial samples | None |
| ACQ42024.1 | Beta-lactamase (blaTEM) |  |  |
| ACQ42045.1 | AAC(6')-Ib-cr |  |  |
| ACQ42046.1 | Beta-lactamase (blaOXA) |  |  |
| ACQ42051.1 | Beta-lactamase (blaCTXM15) |  |  |
| ACQ42056.1 | Dihydrofolate reductase |  |  |
| ACQ42065.1 | Macrolide 2 phosphotransferase<br>(Mph (A)) |  |  |
| ACQ42094.1 | Putative HTH-type<br>transcriptional regulator (YfaX) |  |  |
| ACQ42102.1 | hypothetical protein XCV |  |  |
| ACQ42108.1 | hypothetical protein |  |  |
| ACQ41973.1 | Antitoxin CcdA | cefotaxime-,<br>imipenem-<br>ciprofloxacin-treated<br>bacteria | Control, Ampicillin<br>treated bacteria |
| ACQ41974.1 | Toxin CcdB | Control, ampicillin-,<br>cefotaxime-,<br>ciprofloxacin-treated<br>bacteria | Imipenem |
| ACQ42006.1 | Relaxosome protein TraM | Control, cefotaxime<br>and imipenem<br>treated bacteria | Ampicillin and<br>ciprofloxacin treated<br>bacteria |
| ACQ42109.1 | Uncharacterized protein | Control and<br>imipenem-treated<br>bacteria | Ampicillin,<br>cefotaxime and<br>ciprofloxacin treated<br>bacteria |
| ACQ42036.1 | mRNA interferase (PemK) | Control, ampicillin-,<br>cefotaxime-, | Ciprofloxacin treated<br>bacteria |
| ACQ42069.1 | 34 kDa membrane antigen (Tpd) | imipenem-treated<br>bacteria |  |

The data presented here is from the pre-imputed dataset (Suppl 1) and identifies the protein present in each sample.

**Table 2.**
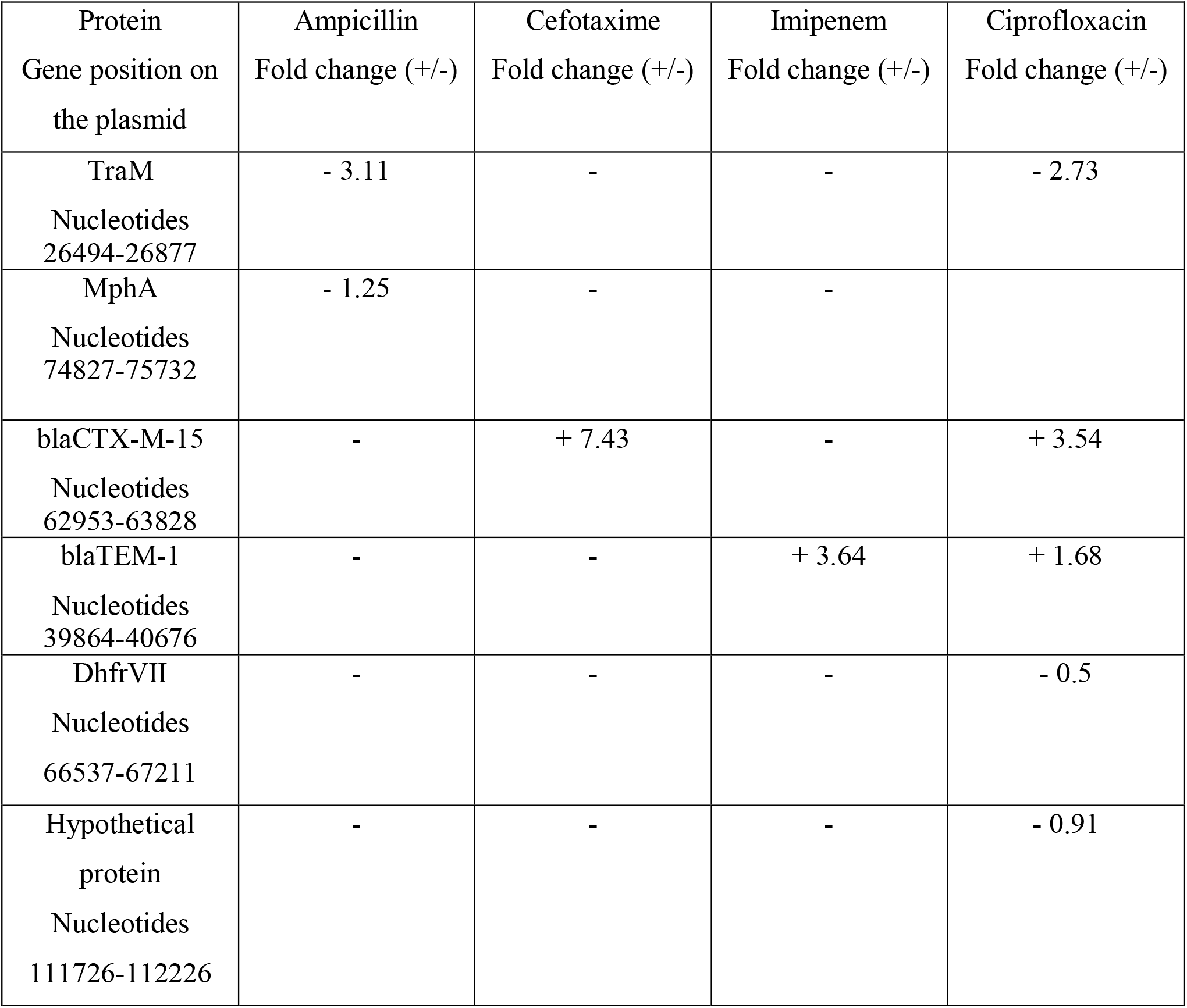
Variations in protein abundance in comparison with control in proteins produced from plasmid genes (post-imputation)

In the presence of ampicillin or ciprofloxacin there was an absence of the protein TraM. TraM is a mating signal, which is used in combination with the integration host factor to bind the *oriT* and prepare the plasmid for transfer. TraM is controlled by an independent promoter to the remainder of the conjugation machinery. Some of the repression systems of TraM include the H-NS repression or the Hfq binding of mRNA transcripts of *traM* or by GroEL chaperone proteins that directly activate proteolysis. The relative abundances of these proteins were not increased in the ampicillin or ciprofloxacin treated *E. coli*. Thus, the lack of TraM was not as a direct result of known repression proteins. The proteins with increased abundance under ampicillin or ciprofloxacin stress relative to the control but absent or with reduced abundance under cefotaxime and imipenem stress comprised 13 proteins (RimO, RfbB, MetK, GalM, RplD, NagZ, RplC, GreA, Apt, SeqA, FumA, SucB and TufB).

The relative increase in the amount of blaCTX-M-15 in the presence of cefotaxime and ciprofloxacin suggest that the production of this protein is regulated, but not only by the direct presence of the cephalosporin alone as ciprofloxacin is a fluoroquinolone. On analysis of the common proteins with increased or decreased abundance across both datasets, no specific protein or pathways were identifiable as potential control systems. The proteins produced in the increased abundances in cefotaxime and ciprofloxacin treated samples within the common proteins across these samples were YjiM (uncharacterised protein), CspA and CspE (cold shock proteins), DeaD (ATP-dependent RNA helicase) and LsrF (terminal protein in the quorum sensing signal autoinducer-2 processing pathway). YjiM and CspE were also identifed in increased abundance in the imipenem treated samples. How the other proteins interact with the plasmid and specifically the blaCTX-M-15 protein production remains to be determined and requires further investigation. The CspA and DeaD proteins are both stress response proteins, but the link to LsrF is unknown. LsrF is produced in the response to the quorum sensing autoinducer AI-2 signal and is thought to promote AI-2 degradation or feedback control to the Lsr operon but has not been associated with antimicrobial resistance (12). The proteins with reduced abundances relative to the control include SpeE (Polyamine aminopropyltransferase), AraA (L-arabinose isomerase), DkgA (2,5-diketo-D-gluconic acid reductase A) and MtlD (Mannitol-1-phosphate 5-dehydrogenase). There was no commonality was identified between these proteins.

blaTEM-1 protein was increased relative to the control in the presence of imipenem or ciprofloxacin but not ampicillin or cefotaxime. This was unexpected as it is a beta-lactamase enzyme and as such if it’s production is controlled we would expect that all ß-lactams induce the ß-lactamase. It also indicates a regulation of ß-lactamase protein production under different antimicrobial stress. The proteins increased in abundance that were unique to the imipenem and ciprofloxacin treated samples were RcsB, ClpX, and GcvP. Of these, RcsB is associated with response to acid stress and ClpX is involved in response to stress. The only common proteins decreased in the largest abundances across ciprofloxacin and imipenem treated samples was YciF, a protein of unknown function.

## Discussion

In this study, quantitative and qualitative proteomics was employed to provide novel insights into the response of plasmid mediated multidrug resistance in *E. coli* to different antimicrobials. While antimicrobials have specific targets on which they exert their mechanism of action, the response of bacteria to these drugs is not limited to the target sites alone (13, 14). Proteomic analysis revealed similarities in the proteome of bacteria in response to antimicrobial-induced stress despite differences in the types of antimicrobials to which they were exposed. Moreover the data presented here highlights the significance of the chromosome-mediated response to antibiotic-induced stress coupled with the resistance mechanism response of the bacteria.

There is limited data on the proteomes of plasmids and their associated bacterial hosts and none that investigate multiple antimicrobial responses in the same pathogen. One study investigated the impact of blaCTX-M-1 *E. coli* to cefotaxime at low and high concentrations and identified that Tra-proteins (including TraM) were significantly upregulated in the presence of high levels (126 mg/L) of cefotaxime but there was no differences at low levels of cefotaxime (0.016 mg/L) (4). Our results do not concur with these findings as we did not identify any significant difference between the cefotaxime treated and the control protein abundances for TraM or the other conjugal plasmid transfer proteins. This study also identified an increase in blaCTX-M-1, PilS and a HEAT domain protein when exposed to 128 mg/L cefotaxime. Our results concur with the increase in blaCTX-M but not the other proteins. However, our results were using an even higher cefotaxime concentration (256 mg/L) and contained blaCTX-M-15 rather than blaCTX-M-1 and a MDR plasmid rather than a single AMR containing plasmid, so this may have influenced for the variation in proteomes. In addition Møller et al., (2017) suggest that the upregulation of the *tra* genes in the presence of cefotaxime was dependent on the presence of blaCTX-M-1 (4). Thus, the difference between blaCTX-M-1 and blaCTX-M-15 may be the reason for the variation. Our study also identified other proteins of potential interest in response to antimicrobial treatment that may aid in the understanding of the control or production of the AMR proteins from the plasmid. A study of carbapenemase producing *E. coli* under carbapenem stress identified increased abundance in GroES in *E. coli* containing blaIMP or blaKPC or blaNDM (in increasing order of abundance) (14). Our study also identified increased GroES in the presence of imipenem, but not the other ß-lactams. However, pEK499 does not contain any carbapenemase and is imipenem susceptible. Thus, we suggest that this is a carbapenem induced response rather than a resistance response, which may be increased by carbapenemase degraded carbapenem as well as whole carbapenem.

Our analysis identified several statistically significant differentially abundant (SSDA) proteins common to groups exposed to the β-lactam antimicrobials but not ciprofloxacin, indicating a β-lactam stress response to exposure from this class of drugs, irrespective of resistance or susceptibility. These include ecotin and free methionine-R-sulfoxide reductase. Ecotin is a serine protease located in the bacterial periplasm and provides the cell with a defence mechanism against host proteases such as neutrophil elastase (15, 16). Free methionine-R-sulfoxide reductase is associated with maintaining redox homeostasis (17, 18). Interestingly, compared to the control, β-lactamase was increased in all antimicrobial-exposed groups except in ampicillin-treated cells where it was not detected at statistically significant levels. In the ampicillin treated samples proteins associated with the outer membrane and cell wall biosynthesis, peptidoglycan-associated lipoprotein (Pal) and beta-hexosaminidase (NagZ), were increased by 4.07-fold and 2.02-fold respectively, indicating an apparent attempt to maintain cell wall integrity during antimicrobial challenge. It has been reported that ampicillin enhances the release of outer membrane vesicles (OMVs) in which Pal is contained, thereby increasing Pal levels (19). This may be of clinical importance because OMVs containing Pal also contain lipopolysaccharides and other inflammatory molecules and the ampicillin-mediated release of OMV from the bacterial cell may contribute to inflammation in the host (19). In this data set, Pal was not detected at statistically significant levels in any other samples. There was a statistically significant decrease in the relative abundance of the plasmid mediated macrolide phosphotransferase (MphA) in ampicillin-treated samples only. A decrease in the levels of MphA indicate that ampicillin may affect the production of this protein, perhaps by activating a repressor, thereby reducing the ability of the bacterial cell to generate resistance. This finding warrants further investigation as it may provide useful information when designing therapeutic regimen involving combination treatments (20). The reduction in the relative abundance of the macrolide 2-phosphotransferase (MphA) occurred in the presence of ampicillin only. There has not previously been any associations between MphA and ampicillin. We did not identify any patterns that could account for repression of MphA production in the presence of ampicillin.

Analysis of the ampicillin treated post-imputation proteomic dataset revealed a significant decrease in the relative abundance of the relaxosome protein TraM, which is responsible for DNA transfer by conjugation between cells (21). Compared to the control, TraM was decreased by 8.61 fold in the ampicillin-treated cells. Mrr restriction system protein, mrr, is involved in the acceptance of foreign DNA from a donor cell (22), and it too was decreased in this group (−2.88 fold decrease). These results suggests fewer plasmid transfer events compared to the control which is in contrast to the finding by Liu et al., (2019) who demonstrated that sub MIC levels of cefotaxime, ampicillin and ciprofloxacin in fact increase the levels of plasmid transfer (23). Differences in plasmids and plasmid-mediated resistance to ampicillin may account for the different findings here. However, the pEK499 plasmid does not confer resistance to ciprofloxacin, but reduced susceptibility, and in this study, the relative abundance of TraM was also decreased in this group indicating lower levels of DNA transfer. Although the levels of TraM are decreased during stationary phase (24), this does not explain the lower levels of TraM in the ciprofloxacin and ampicillin-treated groups compared to the control observed in this dataset. Overproduction of reactive oxygen species (ROS) is known to trigger conjugative transfer (25). In this study the levels of proteins associated with a response to ROS in groups treated with ampicillin or ciprofloxacin were relatively low compared to the control, thus, reduced levels of oxidative stress in these groups may be responsible for a decrease in the relative abundance of proteins associated with gene transfer. Of the β-lactam-exposed groups, the levels of proteins associated with an increase in oxidative stress were greater in groups treated with cefotaxime. For example there was a significant increase in the relative abundance of glutaredoxin 3 GrxC (2.42-fold), glutaredoxin 4 GrxD (1.70-fold), peroxiredoxin OmsC (2.36-fold), thiol peroxidase Tpx (1.80-fold) and peroxide stress resistance protein YaaA (4.06-fold). In contrast, enrichment analysis performed in STRING on SSDA proteins revealed a general decrease in the pathways associated with glycolysis and glyoxylate metabolism. This indicates that the energy used to combat oxidative stress is at the expense of carbohydrate metabolism (26). STRING analysis also revealed an increase in protein levels associated with the ribosome. Because oxidative stress presents bacterial cells with unfavourable environmental conditions, the ability to alter RNA turnover is essential for survival via adaptation to harsh environments. Oxidative stress alters ribosomal activity in bacteria allowing cells to adapt to unfavourable environmental conditions (27). DEAD-box helicases are a group of proteins associated with ensuring continuation of optimal ribosomal activity (28). In addition to the range of proteins associated with translation, the proteomic dataset here identified the protein product of DeaD which increased by 3.02-fold in cefotaxime-treated cells compared to the control. Taken together, comparative proteomic analysis of cefotaxime-exposed cells and untreated cells indicate that cefotaxime induces an oxidative stress response in *E. coli* which is met by an increase in ribosomal activity and a decrease in carbohydrate metabolism.

Imipenem also induced stress in bacterial cells as demonstrated by the number of proteins associated with oxidative stress and their increase in relative abundance compared to the control (e.g. OxyR, Eco, and HdeB). Levels of flagellin protein, FliC, was reduced by almost 22-fold (21.92-fold decrease) in cells exposed to imipenem. This suggests that imipenem induces morphological changes to the bacterial cell which may ultimately affect the motility and adherence properties of the cells. Sub-inhibitory concentrations of antimicrobials are known to induce morphological changes in bacterial cells (29, 30). Understanding these changes and how they may affect bacterial interactions with the host cell are important for developing therapeutic strategies (30). The relative abundance of several proteins involved in carbohydrate metabolism, specifically glucose metabolism, was decreased in this group but the levels of proteins associated with monosaccharide transport into the cell had increased. High affinity transport systems are known to increase under conditions of nutrient limitations (31). It is possible that a decrease in glucose availability induced an increase in the uptake of alternative carbon sources, causing an increase the levels of transporters such as xylose and arabinose. One of the transporters identified in the dataset arising from imipenem-treated bacteria was D-xylose-binding periplasmic protein, encoded by the *xylF* gene. This gene is upregulated in response to cold shock (32). Cold shock inducible genes, while providing protection against temperature decline, also play a role in the bacterial response to antimicrobial stress (32 - 24). In total, there are nine cold shock proteins (Csp) in *E. coli* (CspA-CspI). In this study, the relative abundance of one of these Csp, CspE, was increased in imipenem-, cefotaxime- and ciprofloxacin-treated groups. CspE is constitutively expressed and is responsible for the stability of RNA transcripts arising from genes associated with a general stress response, specifically the master regulator, RpoS (35). The relative abundance of four Csp were increased in bacteria exposed to ciprofloxacin (CspA, CspC, CspD and CspE), suggesting a major role for these proteins in response to ciprofloxacin-induced stress. Compared to the control, there was a 5.58-fold increase in the level of CspD. CspD is generally associated with a carbon-starvation induced stress response during stationary phase growth (36). This protein binds to single stranded DNA and inhibits its replication (37). The significant increase in its abundance compared to the control in this study, indicates that CspD may have a role to play in protection against ciprofloxacin, perhaps by inhibiting DNA replication thereby reducing the effect of ciprofloxacin on this process. Furthermore, there was a 1.41-fold decrease in dihydrofolate reductase, a crucial enzyme for the biosynthesis of DNA precursors. This indicates a reduction in the biosynthesis of DNA in bacterial cells exposed to ciprofloxacin. CpsD is involved in the MqsR/MqsA-mediated toxin/antitoxin (TA) system which regulates the formation of persister cells by inducing biofilm formation (38, 39). Other proteins associated with toxin-anti-toxin system-dependent persister cell formation are Lon, ClpX and Fis, all of which were increased in relative abundance in ciprofloxacin-treated bacteria (38, 39). In addition to CspE, CspA and CspC are single stranded DNA and RNA binding proteins involved in the stabilization of DNA and RNA transcripts under cellular stress and as an adaptation response to low temperatures (40, 41). These proteins increase the half-life of RNA transcripts arising from the expression of stress-induced genes and interfere with the formation of secondary structures in RNA that can result in transcriptional termination (40, 42). It was interesting therefore, to observe an increase in the relative abundance of a substantial number of proteins associated with translation in ciprofloxacin-exposed bacteria compared to the control. The levels of these proteins indicate increased translational activity in this group. In contrast, there was a decrease in the relative abundance of several proteins associated with oxidative stress including superoxide dismutase (SodC), which was reduced by almost 65-fold compared to the control. Taken together the data in this study suggest that exposure to sub inhibitory levels of ciprofloxacin induces a Csp-response which may be, in part, responsible for the increased levels of ribosomal proteins and decrease in proteins associated with oxidative stress. Although ciprofloxacin inhibits DNA replication by targeting DNA topoisomerase and DNA-gyrase, the dataset in this study revealed a significant increase in the relative abundance of plasmid-associated β-lactamase (11.63-fold increase) and of other components involved in cell wall assembly including alanine racemase (Alr) and β-hexosaminidase (NagZ). Compared to the control, the ciprofloxacin-exposed bacteria were the only bacteria with increased abundance of both blaCTX-M-15 and blaTEM-1 β-lactamase, despite cell wall biosynthesis not being the target for the mechanism of action of ciprofloxacin. An increase in the levels of β-lactamase suggests a secondary effect of ciprofloxacin, one which impacts the bacterial cell wall. This observation supports the theory that antimicrobials may serve as an environmental signal for bacteria which induces physiological alterations that provide cells with a competitive advantage (43).

SeqA was one of the 13 proteins with increased abundance under ampicillin or ciprofloxacin stress relative to the control but absent or with reduced abundance under cefotaxime and imipenem stress. SeqA has been identified as a negative modulator of initiation of replication and of plasmid replication (44). We propose that under ampicillin and ciprofloxacin stress SeqA performs this function thus reducing the relative abundance of TraM. However, this does not occur in the presence of cefotaxime or imipenem and is therefore not a general response to antibiotics. As the blaTEM-1 protein and the acid stress response were increased relative to the control in the presence of imipenem or ciprofloxacin but not ampicillin or cefotaxime, we question whether the blaTEM-1 protein production was increased in response to these stress proteins being elevated or to the antimicrobials directly or if the acid stress response is activated in response to the increased blaTEM production. The FruB and YciF proteins present in reduced abundance unique to imipenem and ciprofloxacin have been reported to be upregulated in response to acid stress. Thus, while components of the response to acid stress were increased only some of the proteins required for response and resistance to acid stress were associated with these bacteria.

The response of HdeB and OsmY were opposite in imipenem to ciprofloxacin, i.e. increased in imipenem treated but decreased in ciprofloxacin treated samples. There were no significant changes in the presence of ampicillin or cefotaxime. In the presence of ciprofloxacin but not the other antimicrobials the level of GadB was reduced 7.95-fold relative to the control and in the imipenem treated samples the GadC protein was reduced in abundance 1.84fold. This is interesting to note, as GadBC are usually increased in response to acid stress like the other proteins described. A GadB knockout mutant demonstrated increased persister formation under ampicillin stress (45). In addition, HdeAB, OsmY and OsmE were repressed in persister forming cells (45). The relative protein abundances of HdeB, GadB and OsmY were reduced in the ciprofloxacin treated bacteria, suggesting that these bacteria were persisters. The opposite occurred in the imipenem treated bacteria, as both HdeB and OsmY were increased relative to the control. Hong et al., described both bacterial resistance and persistence in response to stress, such as acid or antimicrobials (45). This proteomics study suggests the specific antimicrobial responses of *E. coli* to these stresses as resistance in relation to ampicillin, cefotaxime and imipenem and persistence in relation to ciprofloxacin. Persistence was demonstrated to occur due to the downregulation of the acid (*gadB, gadX*), osmotic (*osmY*), and multidrug (*mdtF*) resistance systems due to the degradation of MqsA by proteases (ClpXP and Lon) (45). Using the proteomics data we identified that only in the presence of ciprofloxacin were the Lon and ClpX proteins increased in abundance together with decreases in abundance of the GadB, HdeB and OsmY proteins. This pattern is described in the persister formation rather than the resistance formation induced pathways. In addition, CpsD is involved in the MqsR/MqsA-mediated toxin/antitoxin (TA) system which regulates the formation of persister cells by inducing biofilm formation (38, 39). While we detected increased CspD only in ciprofloxacin-treated bacteria, we did not detect MqsR or MsqA proteins in any sample.

## Conclusions

The data presented in this study has provided novel insights into the changes that occur in the proteome of multidrug resistant *E. coli* when challenged with different antimicrobials, and highlight a significant role for chromosomally-encoded genes in the response of bacteria to these antimicrobials. The data arising from proteomic analysis of *E. coli* challenged with three different β-lactam antibiotics identified distinct differences in the cellular response to each drug. These data also identified novel proteins not previously associated with AMR or antimicrobials responses in pathogens.

## Acknowledgements

The mass spectrometry facilities were funded by a Science Foundation Ireland infrastructure award to SD (SFI 12/RI/2346(3)).

## References

1. https://www.who.int/medicines/publications/WHO-PPL-Short_Summary_25Feb-ET_NM_WHO.pdf (accessed 14/07/2021)

2. Woodford N, Carattoli A, Karisik E, Underwood A, Ellington MJ, Livermore DM. Complete nucleotide sequences of plasmids pEK204, pEK499, and pEK516, encoding CTX-M enzymes in three major Escherichia coli lineages from the United Kingdom, all belonging to the international O25:H4-ST131 clone. Antimicrob Agents Chemother. 2009 Oct;53(10):4472–82.

3. Blumenscheit C, Pfeifer Y, Werner G, John C, Schneider A, Lasch P, Doellinger J. Unbiased antimicrobial resistance detection from clinical bacterial isolates using proteomics. bioRxiv 2020.11.17.386540;

4. Møller TSB, Liu G, Boysen A, Thomsen LE, Lüthje FL, Mortensen S, Møller-Jensen J, Olsen JE. Treatment with Cefotaxime Affects Expression of Conjugation Associated Proteins and Conjugation Transfer Frequency of an IncI1 Plasmid in Escherichia coli. Front Microbiol. 2017 Nov 29;8:2365.

5. Møller TSB, Liu G, Hartman HB, Rau MH, Mortensen S, Thamsborg K, Johansen AE, Sommer MOA, Guardabassi L, Poolman MG, Olsen JE. Global responses to oxytetracycline treatment in tetracycline-resistant Escherichia coli. Sci Rep. 2020 May 21;10(1):8438.

6. Hubner NC, Bird AW, Cox J, Splettstoesser B, Bandilla P, Poser I, Hyman A, Mann M. Quantitative proteomics combined with BAC TransgeneOmics reveals in vivo protein interactions. J Cell Biol. 2010 May 17;189(4):739–54.

7. Côté RG, Griss J, Dianes JA, Wang R, Wright JC, van den Toorn HW, van Breukelen B, Heck AJ, Hulstaert N, Martens L, Reisinger F, Csordas A, Ovelleiro D, Perez-Rivevol Y, Barsnes H, Hermjakob H, Vizcaíno JA. The PRoteomics IDEntification (PRIDE) Converter 2 framework: an improved suite of tools to facilitate data submission to the PRIDE database and the ProteomeXchange consortium. Mol Cell Proteomics. 2012 Dec;11(12):1682–9.

8. Lesniak J, Barton WA, Nikolov DB. Structural and functional features of the Escherichia coli hydroperoxide resistance protein OsmC. Protein Sci. 2003 Dec;12(12):2838–43.

9. Molina-Heredia FP, Houée-Levin C, Berthomieu C, Touati D, Tremey E, Favaudon V, Adam V, Nivière V. Detoxification of superoxide without production of H2O2: antioxidant activity of superoxide reductase complexed with ferrocyanide. Proc Natl Acad Sci U S A. 2006 Oct 3;103(40):14750–5.

10. Prieto-Alamo MJ, Jurado J, Gallardo-Madueno R, Monje-Casas F, Holmgren A, Pueyo C. Transcriptional regulation of glutaredoxin and thioredoxin pathways and related enzymes in response to oxidative stress. J Biol Chem. 2000 May 5;275(18):13398–405.

11. Smirnova G, Muzyka N, Lepekhina E, Oktyabrsky O. Roles of the glutathione-and thioredoxin-dependent systems in the Escherichia coli responses to ciprofloxacin and ampicillin. Arch Microbiol. 2016 Nov;198(9):913–21.

12. Wang L, Hashimoto Y, Tsao CY, Valdes JJ, Bentley WE. Cyclic AMP (cAMP) and cAMP receptor protein influence both synthesis and uptake of extracellular autoinducer 2 in Escherichia coli. J Bacteriol. 2005 Mar;187(6):2066–76.

13. Kapoor G, Saigal S, Elongavan A. Action and resistance mechanisms of antibiotics: A guide for clinicians. J Anaesthesiol Clin Pharmacol. 2017 Jul-Sep;33(3):300–305.

14. Sidjabat HE, Gien J, Kvaskoff D, Ashman K, Vaswani K, Reed S, McGeary RP, Paterson DL, Bordin A, Schenk G. The use of SWATH to analyse the dynamic changes of bacterial proteome of carbapanemase-producing Escherichia coli under antibiotic pressure. Sci Rep. 2018 Mar 1;8(1):3871.

15. Eggers CT, Wang SX, Fletterick RJ, Craik CS. The role of ecotin dimerization in protease inhibition. J Mol Biol. 2001 May 18;308(5):975–91.

16. Eggers CT, Murray IA, Delmar VA, Day AG, Craik CS. The periplasmic serine protease inhibitor ecotin protects bacteria against neutrophil elastase. Biochem J. 2004 Apr 1;379(Pt 1):107–18.

17. Lin Z, Johnson LC, Weissbach H, Brot N, Lively MO, Lowther WT. Free methionine-(R)-sulfoxide reductase from Escherichia coli reveals a new GAF domain function. Proc Natl Acad Sci U S A. 2007 Jun 5;104(23):9597–602.

18. Wang Z, Xia X, Zhang M, Fang J, Li Y, Zhang M. Purification and Characterization of Glutathione Binding Protein GsiB from Escherichia coli. Biomed Res Int. 2018 Nov 1;2018:3429569.

19. Michel LV, Gallardo L, Konovalova A, Bauer M, Jackson N, Zavorin M, McNamara C, Pierce J, Cheng S, Snyder E, Hellman J, Pichichero ME. Ampicillin triggers the release of Pal in toxic vesicles from Escherichia coli. Int J Antimicrob Agents. 2020 Dec;56(6):106163.

20. Tamma PD, Cosgrove SE, Maragakis LL. Combination therapy for treatment of infections with gram-negative bacteria. Clin Microbiol Rev. 2012 Jul;25(3):450–70. doi: 10.1128/CMR.05041-11.

21. Pölzleitner E, Zechner EL, Renner W, Fratte R, Jauk B, Högenauer G, Koraimann G. TraM of plasmid R1 controls transfer gene expression as an integrated control element in a complex regulatory network. Mol Microbiol. 1997 Aug;25(3):495–507.

22. Waite-Rees PA, Keating CJ, Moran LS, Slatko BE, Hornstra LJ, Benner JS. Characterization and expression of the Escherichia coli Mrr restriction system. J Bacteriol. 1991 Aug;173(16):5207–19.

23. Liu G, Bogaj K, Bortolaia V, Olsen JE, Thomsen LE. Antibiotic-Induced, Increased Conjugative Transfer Is Common to Diverse Naturally Occurring ESBL Plasmids in Escherichia coli. Front Microbiol. 2019 Sep 10;10:2119.

24. Zatyka M, Thomas CM. Control of genes for conjugative transfer of plasmids and other mobile elements. FEMS Microbiol Rev. 1998;21:291–319.

25. Zhang S, Wang Y, Song H, Lu J, Yuan Z, Guo J. Copper nanoparticles and copper ions promote horizontal transfer of plasmid-mediated multi-antibiotic resistance genes across bacterial genera. Environ Int. 2019 Aug;129:478–487.

26. Mullarky E, Cantley LC. Diverting Glycolysis to Combat Oxidative Stress. In: Nakao K, Minato N, Uemoto S, editors. Innovative Medicine: Basic Research and Development [Internet]. Tokyo: Springer; 2015. pp. 3–23.

27. Leiva LE, Pincheira A, Elgamal S, Kienast SD, Bravo V, Leufken J, Gutiérrez D, Leidel SA, Ibba M, Katz A. Modulation of Escherichia coli Translation by the Specific Inactivation of tRNAGly Under Oxidative Stress. Front Genet. 2020 Aug 18;11:856.

28. Redder P, Hausmann S, Khemici V, Yasrebi H, Linder P. Bacterial versatility requires DEAD-box RNA helicases. FEMS Microbiol Rev. 2015 May;39(3):392–412.

29. Lorian V. Medical relevance of low concentrations of antibiotics. J Antimicrob Chemother. 1993 May;31 Suppl D:137–48. doi: 10.1093/jac/31.suppl_d.137.

30. Fonseca AP, Sousa JC. Effect of antibiotic-induced morphological changes on surface properties, motility and adhesion of nosocomial Pseudomonas aeruginosa strains under different physiological states. J Appl Microbiol. 2007 Nov;103(5):1828–37.

31. Ferenci T. Adaptation to life at micromolar nutrient levels: the regulation of Escherichia coli glucose transport by endoinduction and cAMP. FEMS Microbiol Rev. 1996 Jul;18(4):301–17.

32. Phadtare S, Inouye M. Genome-wide transcriptional analysis of the cold shock response in wild-type and cold-sensitive, quadruple-csp-deletion strains of Escherichia coli. J Bacteriol. 2004 Oct;186(20):7007–14.

33. Xia B, Ke H, Inouye M. Acquirement of cold sensitivity by quadruple deletion of the cspA family and its suppression by PNPase S1 domain in Escherichia coli. Mol Microbiol. 2001 Apr;40(1):179–88.

34. Cruz-Loya M, Kang TM, Lozano NA, Watanabe R, Tekin E, Damoiseaux R, Savage VM, Yeh PJ. Stressor interaction networks suggest antibiotic resistance co-opted from stress responses to temperature. ISME J. 2019 Jan;13(1):12–23.

35. Shenhar Y, Biran D, Ron EZ. Resistance to environmental stress requires the RNA chaperones CspC and CspE. Environ Microbiol Rep. 2012 Oct;4(5):532–9.

36. Yamanaka K, Inouye M. Growth-phase-dependent expression of cspD, encoding a member of the CspA family in Escherichia coli. J Bacteriol. 1997 Aug;179(16):5126–30.

37. Yamanaka K, Zheng W, Crooke E, Wang YH, Inouye M. CspD, a novel DNA replication inhibitor induced during the stationary phase in Escherichia coli. Mol Microbiol. 2001 Mar;39(6):1572–84.

38. Kim Y, Wood TK. Toxins Hha and CspD and small RNA regulator Hfq are involved in persister cell formation through MqsR in Escherichia coli. Biochem Biophys Res Commun. 2010 Jan 1;391(1):209–13.

39. Kim Y, Wang X, Zhang XS, Grigoriu S, Page R, Peti W, Wood TK. Escherichia coli toxin/antitoxin pair MqsR/MqsA regulate toxin CspD. Environ Microbiol. 2010 May;12(5):1105–21.

40. Bae W, Xia B, Inouye M, Severinov K. Escherichia coli CspA-family RNA chaperones are transcription antiterminators. Proc Natl Acad Sci U S A. 2000 Jul 5;97(14):7784–9.

41. Phadtare S, Tadigotla V, Shin WH, Sengupta A, Severinov K. Analysis of Escherichia coli global gene expression profiles in response to overexpression and deletion of CspC and CspE. J Bacteriol. 2006 Apr;188(7):2521–7.

42. Phadtare S, Inouye M. Role of CspC and CspE in regulation of expression of RpoS and UspA, the stress response proteins in Escherichia coli. J Bacteriol. 2001 Feb;183(4):1205–14.

43. Linares JF, Gustafsson I, Baquero F, Martinez JL. Antibiotics as intermicrobial signaling agents instead of weapons. Proc Natl Acad Sci U S A. 2006 Dec 19;103(51):19484–9.

44. Douraid D, Ahmed L. SeqA, the Escherichia coli origin sequestration protein, can regulate the replication of the pBR322 plasmid. Plasmid. 2011 Jan;65(1):15–9.

45. Hong SH, Wang X, O’Connor HF, Benedik MJ, Wood TK. Bacterial persistence increases as environmental fitness decreases. Microb Biotechnol. 2012 Jul;5(4):509–22.

