## Supplemental Figures 1 to 4 for "Global protein responses of multi-drug resistant plasmid containing *Escherichia coli* to ampicillin, cefotaxime, imipenem and ciprofloxacin"

### Supplementary Figures 1 – 4

**Fig. 1.** STRING pathway analysis depicting interactions between stat. sig. proteins increased (A) or decreased (B) in relative abundance arising from pairwise t-tests performed on proteins detected in ampicillin-treated groups and controls.

**Fig. 2.** STRING pathway analysis depicting interactions between stat. sig. proteins increased (A) or decreased (B) in relative abundance arising from pairwise t-tests performed on proteins detected in cefotaxime-treated groups and controls.

**Fig. 3.** STRING pathway analysis depicting interactions between stat. sig. proteins increased (A) or decreased (B) in relative abundance arising from pairwise t-tests performed on proteins detected in imipenem-treated groups and controls.

**Fig. 4.** STRING pathway analysis depicting interactions between stat. sig. proteins increased (A) or decreased (B) in relative abundance arising from pairwise t-tests performed on proteins detected in ciprofloxacin-treated groups and controls.

**Stat. sig.: statistically significant**

**Ampicillin:** STRING including stat. sig. proteins (increased) STRING containing stat. sig. proteins (decreased)

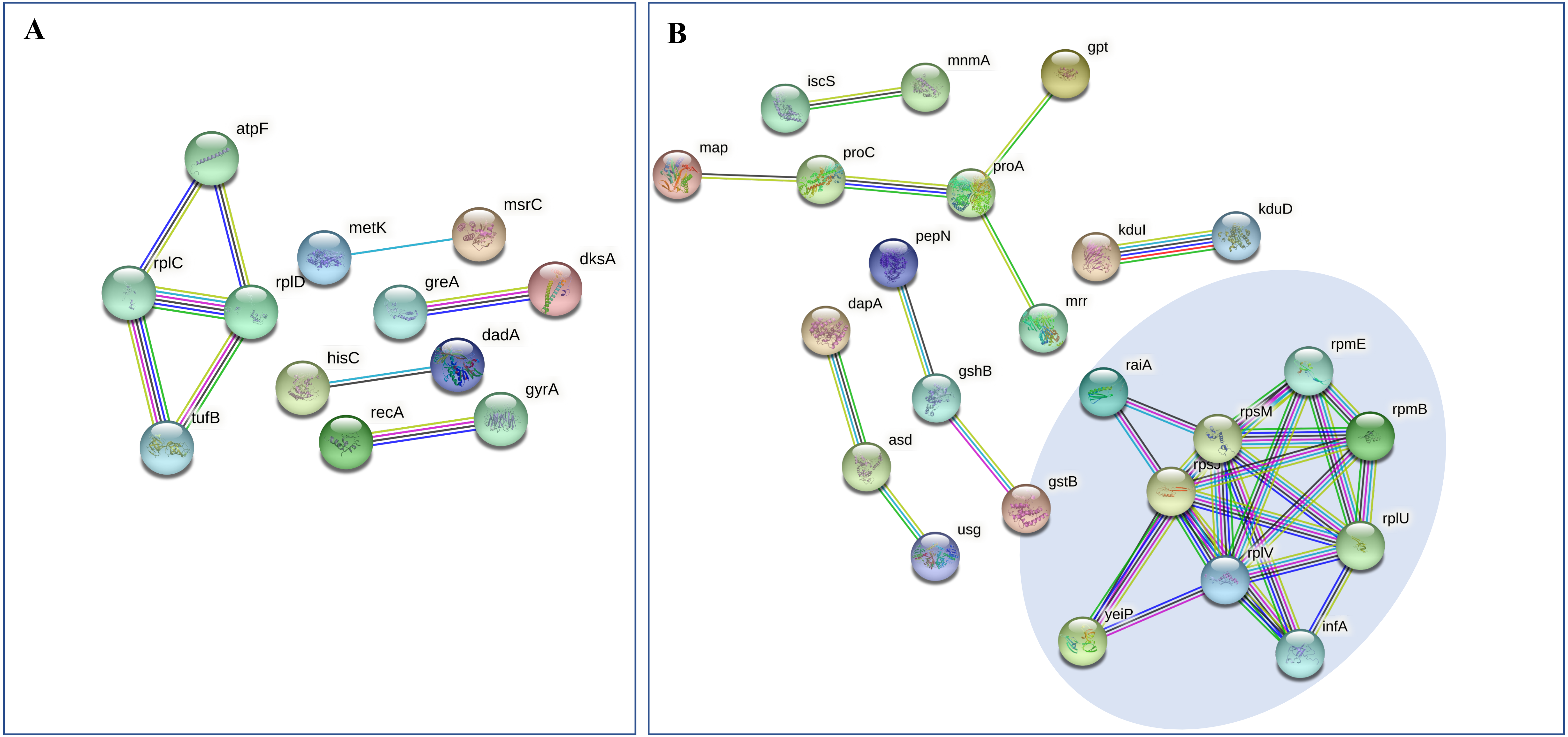


**Supplementary figure 1**: STRING pathway analysis depicts interactions occurring between statistically significant proteins arising from pairwise t-tests between ampicillin-treated bacteria and the control. A decrease in the relative abundance of proteins involved in translation and the ribosome is highlighted in the blue circle (B). STRING interaction score: high confidence (0.700).

**Cefotaxime:**

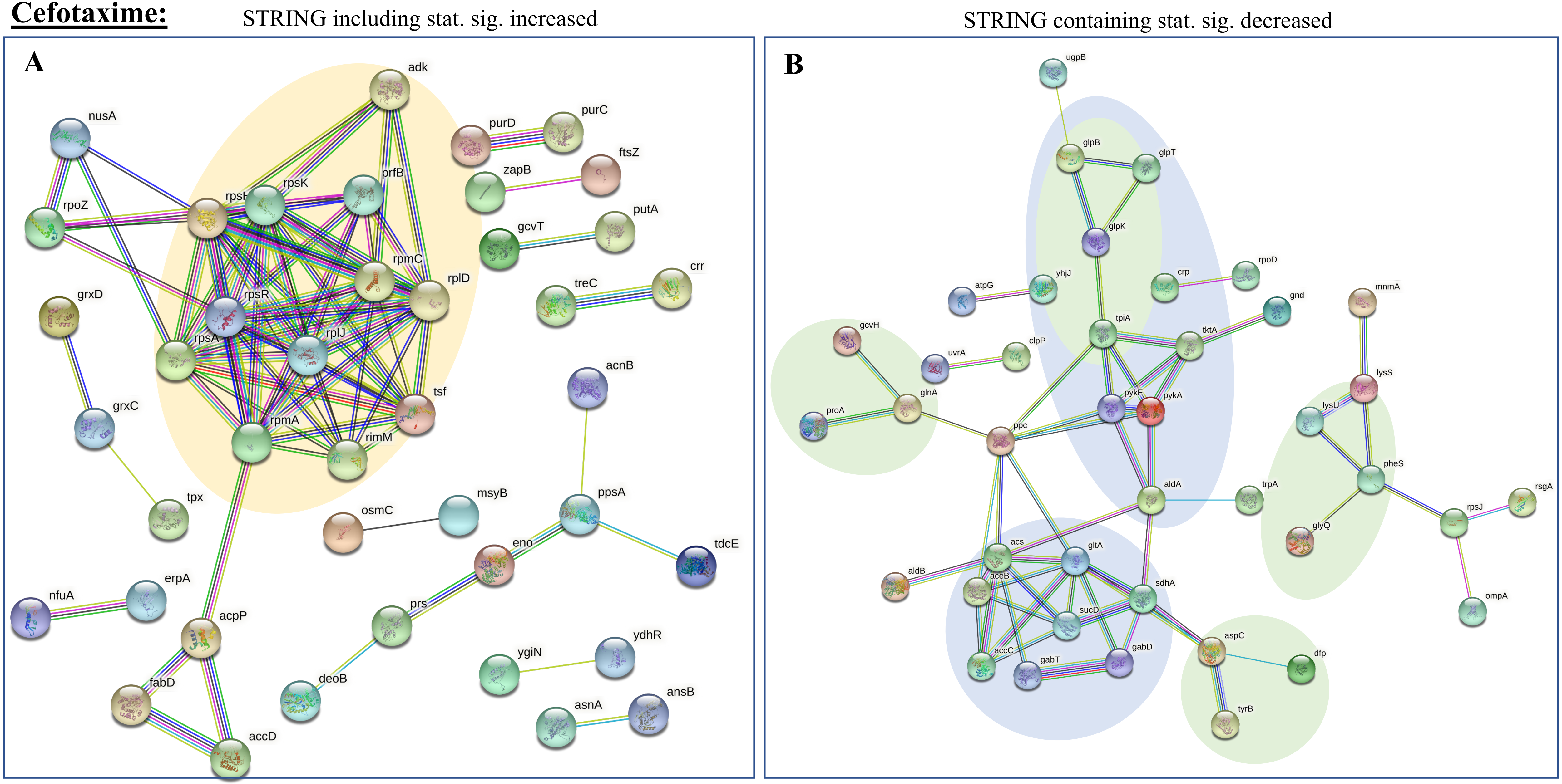


**Supplementary figure 2**: STRING pathway analysis depicts interactions occurring between statistically significant proteins arising from pairwise t-tests between cefotaxime-treated bacteria and the control. An increase in the relative abundance of proteins involved in translation and the ribosome are highlighted in the orange circle (A). A decrease in the relative abundance of proteins associated with carbon metabolism and amino acid metabolism are highlighted in the blue and green circles respectively (B). STRING interaction score: high confidence (0.700).

**Imipenem:** STRING including stat. sig. proteins (increased)

**A**

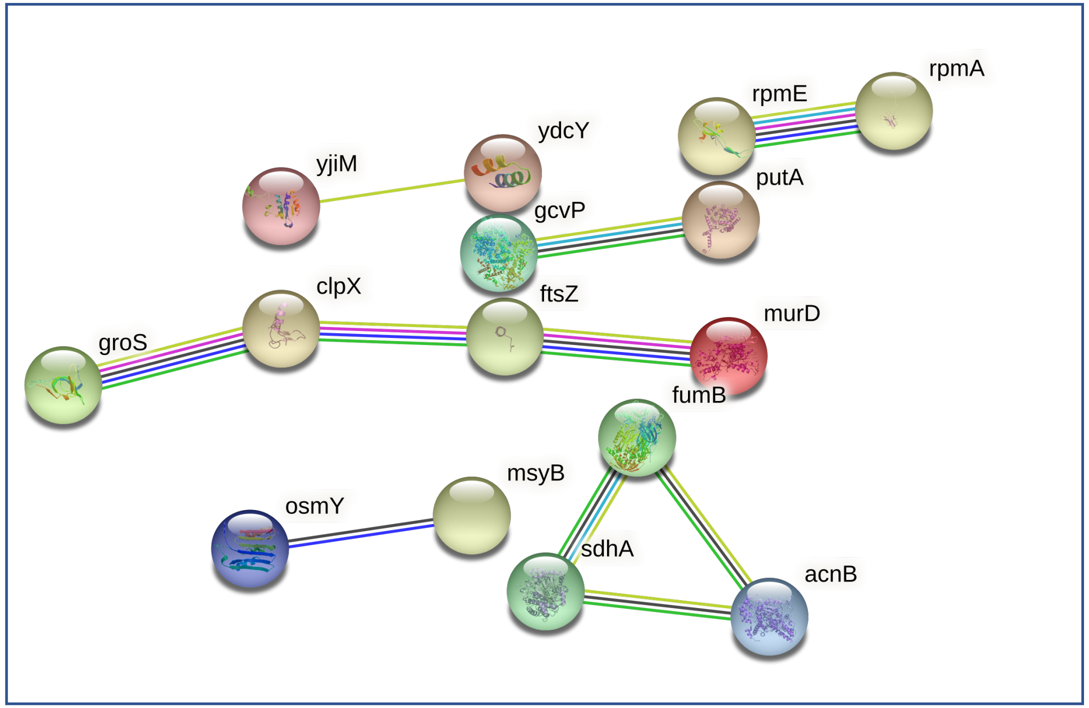


STRING containing stat. sig. (decreased)

**B**

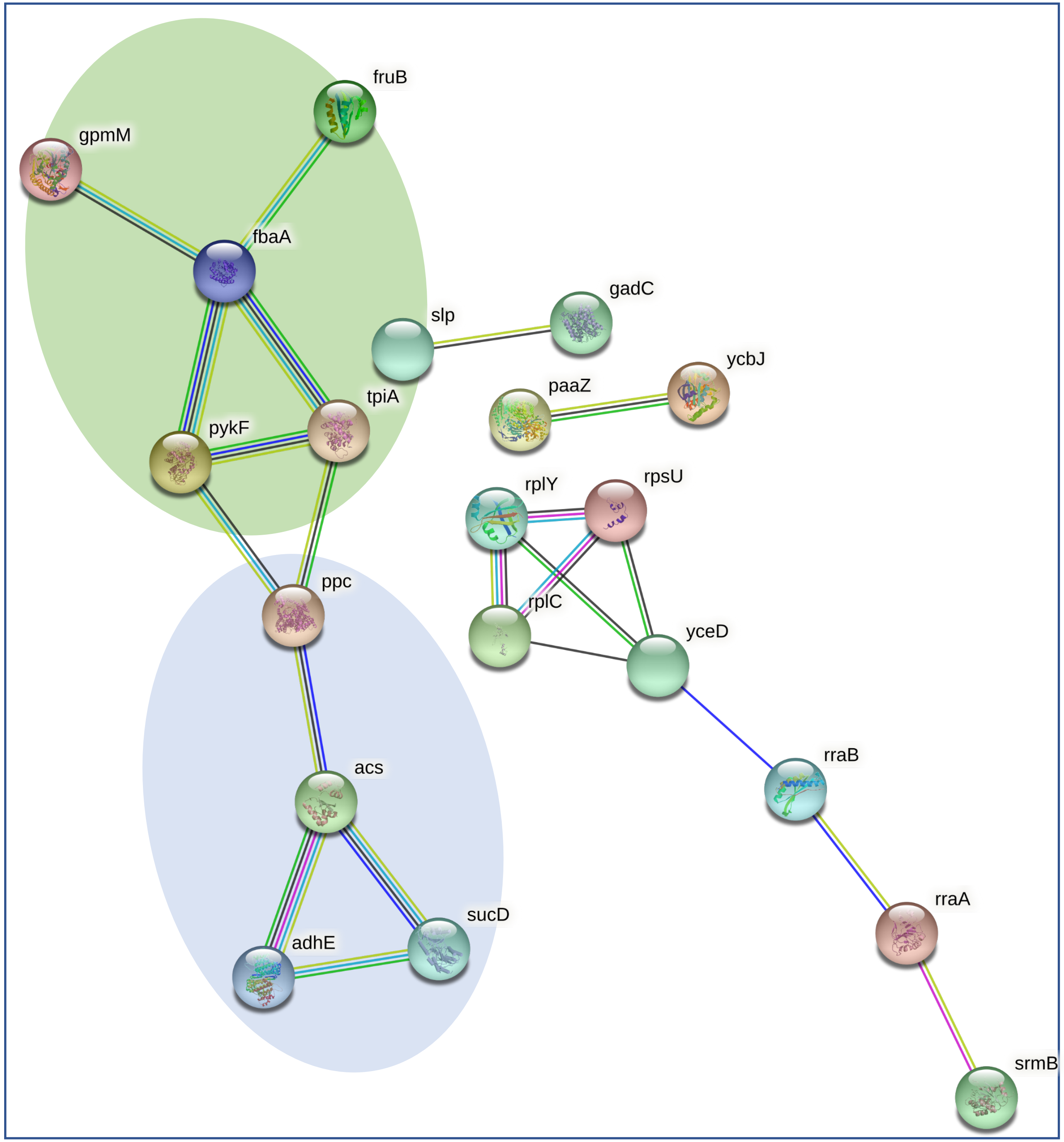


**Supplementary figure 3**: STRING pathway analysis depicts interactions occurring between statistically significant proteins arising from pairwise t-tests between imipenem-treated bacteria and the control. A decrease in the relative abundance of proteins associated with carbon and pyruvate metabolism, and glycolysis are highlighted in the blue and green circles respectively (B). STRING interaction score: high confidence (0.700).

**Ciprofloxacin** STRING including stat. sig. proteins (increased) STRING containing stat. sig. proteins (decreased)

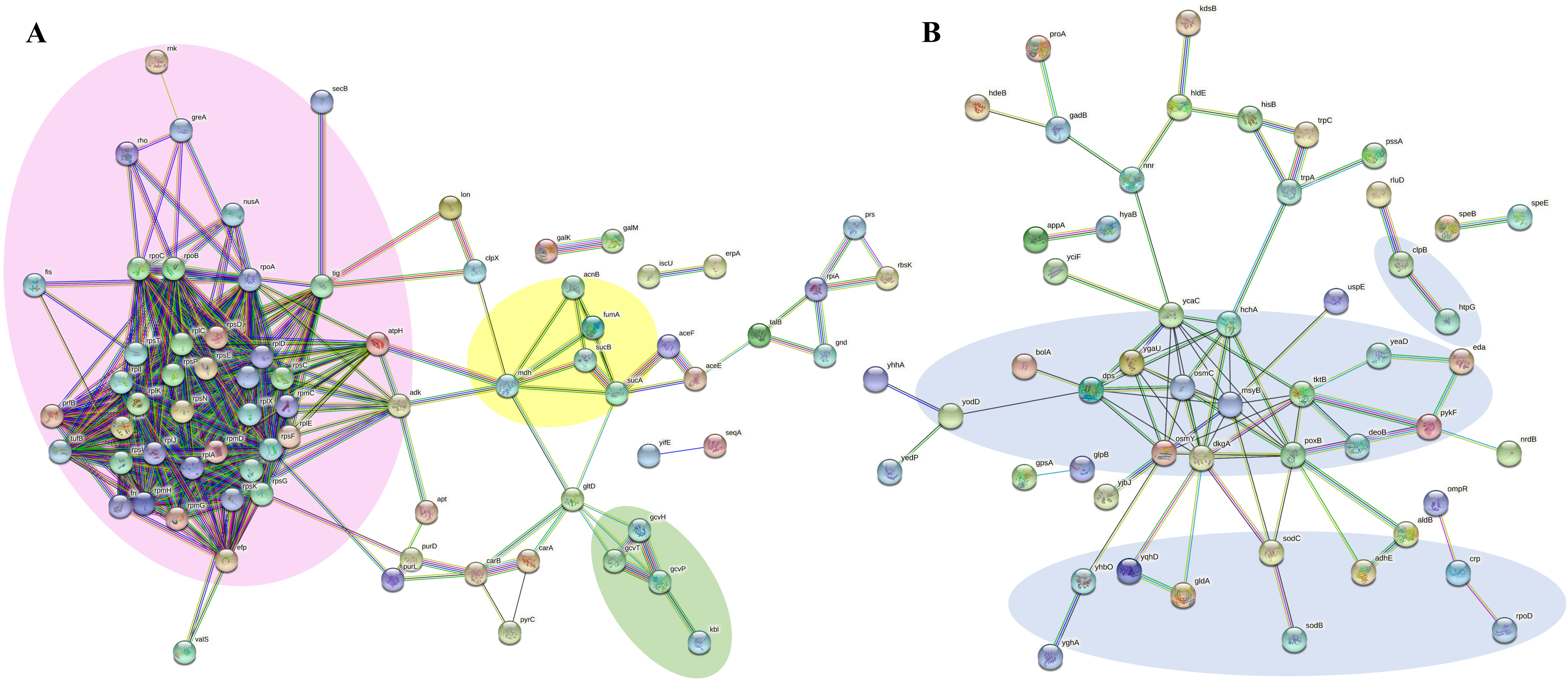


**Supplementary figure 4**: STRING pathway analysis depicts interactions occurring between statistically significant proteins arising from pairwise t-tests between ciprofloxacin-treated bacteria and the control. An increase in the relative abundance of proteins involved in translation and the ribosome, the TCA cycle, and serine, threonine and glycine metabolism, are highlighted in the purple, yellow and green circle respectively (A). A decrease in the relative abundance of proteins associated with a stress response, including oxidative stress, are highlighted in the blue circles (B). STRING interaction score: high confidence (0.700).
